# The Effects of Nitric Oxide on Choroidal Gene Expression

**DOI:** 10.1101/2023.06.16.545343

**Authors:** Makenzie B. Merkley, Diana Soriano, Kenneth L. Jones, Jody A. Summers

**Affiliations:** Department of Biology, University of Oklahoma, Norman, Oklahoma, 73019, United States; Department of Cell Biology, University of Oklahoma Health Sciences Center, Oklahoma City, Oklahoma, 73104, United States; Bioinformatic Solutions, LLC, Seridan Wyoming 82801

## Abstract

**Purpose:** Nitric oxide (NO) is recognized as an important biological mediator that controls several physiological functions, and evidence is now emerging that this molecule may play a significant role in the postnatal control of ocular growth and myopia development. We therefore sought to understand the role that nitric oxide plays in visually-guided ocular growth in order to gain insight into the underlying mechanisms of this process.

**Methods:** Choroids were incubated in organ culture in the presence of the NO donor, PAPA-NONOate (1.5 mM). Following RNA extraction, bulk RNA-seq was used to quantify and compare choroidal gene expression in the presence and absence of PAPA-NONOate. We used bioinformatics to identify enriched canonical pathways, predicted diseases and functions, and regulatory effects of NO in the choroid.

**Results:** Upon treatment of normal chick choroids with the NO donor, PAPA-NONOate, we identified a total of 837 differentially expressed genes (259 upregulated genes, 578 down-regulated genes) compared with untreated controls. Among these, the top five upregulated genes were LSMEM1, STEAP4, HSPB9, and CCL19, and the top five down-regulated genes were CDCA3, SMC2, a novel gene (ENSALGALG00000050836), an uncharacterized gene (LOC107054158), and SPAG5. Bioinformatics predicted that NO treatment will activate pathways involved in cell and organismal death, necrosis, and cardiovascular system development, and inhibit pathways involved in cell proliferation, cell movement, and gene expression.

**Conclusions:** The findings reported herein may provide insight into possible effects of NO in the choroid during visually regulated eye growth, and help to identify targeted therapies for the treatment of myopia and other ocular diseases.

## INTRODUCTION

Nitric oxide (NO) is a short-lived, soluble, free radical gas produced by a wide variety of cells and is acknowledged as a vital intercellular messenger in multiple systems in the body. It has been shown to have numerous functions, one of which is vasodilation [1]. The therapeutic applications of nitric oxide in medicine have been studied for decades, especially in cardiovascular and nervous systems, and in its role in immunological responses [2]. Evidence is now emerging that NO is a mediator of physiological, and possibly pathological, processes in the eye [3–5]. NO regulates ocular hemodynamics and protects vascular endothelial and nerve cells against pathogenic factors associated with glaucoma, ischemia, and diabetes mellitus [4]. NO, constitutively formed by endothelial NOS (eNOS, NOS3) and neuronal NOS (nNOS, NOS1), contributes to vasodilation, increased blood flow, and decreased vascular resistance in ocular circulation. Immunological NOS (iNOS, NOS2) is inducible only in pathological conditions and, once induced, produces large amounts of NO for long periods of time. NO is rapidly converted into nitrite (NO_2_-), nitrate (NO_3_-), peroxynitrite (ONOO^-^), and free radicals to induce pathophysiological actions which can lead to glaucoma, retinopathy, age-related macular degeneration (AMD), and cataracts [5]. Both underproduction and overproduction of NO have been shown to cause various eye diseases and, because of these dichotomous roles, difficulties arise in constructing therapeutic strategies with NO supplementation or NOS inhibition. Underproduction of NO that occurs in conditions such as glaucoma can be partially corrected by providing NOS substrates or NO donors. Conversely, excess NO production that is seen in conditions like diabetic retinopathy, age-related macular degeneration, and ocular inflammation could be corrected by providing NOS inhibitors [5].

Of particular interest is the role that nitric oxide plays in visually guided eye growth. Emmetropization is a visually guided developmental process by which the eye grows in a way to minimize refractive error and maintain emmetropia [6]. While the mechanisms that coordinate this process are not fully understood, results from clinical and experimental studies indicate that emmetropization requires visual input and occurs via a local (within the eye) retina-to-scleral chemical cascade [7, 8].

Myopia (near-sightedness) is a common abnormal visual condition in which objects at infinity are focused in front of the photoreceptors, causing blurred distance vision. Once viewed as a benign refractive condition, myopia has now been shown to be a significant risk factor for blinding eye diseases including glaucoma, retinal detachment, and macular generation, therefore representing a leading cause of blindness worldwide [9, 10]. Although the pathophysiology of myopia development is complex, it is generally accepted that myopia results from a failure of the emmetropization process [7].

Myopia can be induced in animals by depriving the retina of form vision. Deprivation of form vision through the use of visual occluders has been shown to accelerate ocular growth and the development of myopia in vertebrate species including fish, chicks, mice, tree shrews, guinea pigs, and primates [11–15]. Following a period of form deprivation-induced myopia, removal of the occluder results in myopic defocus in the previously form-deprived eye, leading to a rapid deceleration in ocular elongation and return to emmetropia (“recovery”) [16].

The existence of biochemical signaling cascades which induce molecular and structural changes in the retina, choroid and sclera, ultimately leading to altered eye growth and changes in refractive state has been well-established [17]. The choroid in particular has been shown to undergo changes in thickness, permeability, and blood flow through periods of visually guided eye growth [18–22]. While the molecular mechanisms behind these choroidal changes are not yet known, there is increasing evidence that NO may be involved. Administration of the nitric oxide synthase inhibitor, N^a^-nitro-L-arginine methyl ester (L-NAME), has been shown to inhibit the choroidal “recovery” response, resulting in a continued increase in the rate of ocular elongation and continued myopia development [23, 24]. Furthermore, intravitreal delivery of the NO donor, sodium nitroprusside (SNP) or the NOS substrate, L-arginine, has been shown to slow the rate of axial elongation and inhibit the development of myopic refractive error in a dose-dependent manner [25]. Taken together, these results suggest that NO acts to slow myopic eye growth and inhibition of NO synthesis prevents recovery from induced myopia. While neither the cellular source of NO nor the target of NO mediating these effects on eye growth have been identified, we predict that one or more cell population in the choroid is a potential target of NO. Our prediction is based on the observation that administration of L-NAME reduces choroidal concentrations of nitrate [24] and that choroidal expression of interleukin 6 (IL6) is mediated by NO [26]. Therefore, the objective of the current study is to gain an understanding of the biological effects of NO on the chick choroid, by analyzing changes in global gene expression patterns following treatment of choroids with the NO donor, PAPA NONOate.

## METHODS

### Ethics and Animals

Animals were managed in accordance with the ARVO Statement for the Use of Animals in Ophthalmic and Vision Research, with the Animal Welfare Act, and with the National Institutes of Health Guidelines. All procedures were approved by the Institutional Animal Care and Use Committee of the University of Oklahoma Health Sciences Center (protocol # 20-092-H). White Leghorn male chicks (*Gallus gallus*) were obtained as 2-day-old hatchlings from Ideal Breeding Poultry Farms (Cameron, TX). Chicks were housed in temperature-controlled brooders with a 12-hour light/dark cycle and were given food and water ad libitum. At the end of experiments, chicks were euthanized by overdose of isoflurane inhalant anesthetic (IsoThesia; Vetus Animal Health, Rockville Center, NY), followed by decapitation.

### Visual Manipulations

Form deprivation myopia (FDM) was induced in 3 to 4 day-old chicks by applying translucent plastic goggles to one eye, as previously described [27]. The contralateral eyes (left eyes) of all chicks remained untreated and served as controls. Chicks were checked daily for the condition of the goggles. Goggles remained in place for 10 days, after which time the goggles were removed and chicks were allowed to experience unrestricted vision (recover) for 3 hrs, at which time chicks were sacrificed and ocular tissues harvested. A separate group of chicks remained ungoggled for and sacrificed at 15 days of age for organ culture with PAPA-NONOate (see below).

### Tissue Preparation

Chicks were euthanized by an overdose of isoflurane inhalant anesthetic (IsoThesia; Vetus Animal Health). Eyes were enucleated and cut along the equator to separate the anterior segment and posterior eye cup. Anterior tissues were discarded, and the vitreous body was removed from the posterior eye cups. An 8 mm punch was taken from the posterior pole of the chick eye using a dermal biopsy punch (Miltex Inc., York, PA). Punches were located nasal to the exit of the optic nerve, with care to exclude the optic nerve and pecten oculi. With the aid of a dissecting microscope, the retina and majority of RPE were removed from the underlying choroid and sclera with a drop of phosphate buffered saline (PBS; 3 mM dibasic sodium phosphate, 1.5 mM monobasic sodium phosphate, 150 mM NaCl, pH 7.2) and gentle brushing. For DAF-FM diacetate (DAF-2DA) labeling, choroids with sclera still attached were placed into a 48-well flat bottom plate (Corning Inc., Corning, NY). A small amount of RPE was left on the choroids to discriminate between the RPE and scleral side of the tissue. For RNA-seq and Taqman analyses, all RPE was removed from the choroid with gentle brushing and rinsing with PBS, and choroids were placed in wells of a 48 well plate containing culture medium [1:1 mixture of Dulbecco’s Modified Eagle’s Medium and Ham’s F12 containing streptomycin (0.1 mg/ml), penicillin (100 units/ml) and gentamicin (50 μg/ml)].

### NO imaging

NO imaging was performed as reported previously [28, 29]. Chicks were allowed to recover from form deprivation for 3 hours, at which time they were killed and their eyes enucleated. Under dim red light, eyes were hemisected and gently washed with Ringer solution [102 mM NaCl, 28 mM NaHCO_3_, 1 mM CaCl_2_, 1 mM MgCl_2_, 2.6 mM KCl, and 5 mM glucose, equilibrated with 95% O_2_, 5% CO_2_ (pH 7.4)]. An 8 mm punch was obtained from the posterior pole of the eye and the retina, with some adherent RPE was gently removed from the underlying choroid and sclera. An ≈ 8mm diameter piece of nitrocellulose (Schleicher & Schuell, Keene, NH) was then placed onto the choroid and the filter paper, together with the choroid, was peeled away from the underlying sclera. The choroid (scleral side up) and the underlying filter paper were sectioned into ≈200 μm slices using a Tissue Chopper (Narishige Scientific Instrument Lab., Tokyo, Japan). The slices, attached to filter paper, were submerged in cold Ringer solution. The slices were then incubated in darkness in Ringer solution containing the NO-sensitive dye 4,5-diaminofluorescein diacetate (DAF-2DA; 20 μm) for 1 hour at room temperature, while continuously bubbling with room air. Following dye loading, the slices were washed three times with fresh, aerated Ringer solution for 15 min/wash at room temperature, followed by fixation in 4% paraformaldehyde overnight at 4°C. Following fixation, DAF-2DA-labelled choroid slices were washed in PBS (3 x 15 min) at room temperature, then incubated for 10 s at RT with 0.0005% DAPI nuclear stain, followed by a final rinse in PBS. Choroid sections were mounted onto the slides with Prolong Gold Antifade reagent containing DAPI (ThermoFisher Scientific), and the DAF-2DA labeled sections were examined under an Olympus Fluoview 1000 laser-scanning confocal microscope (Center Valley, PA).

### Organ Culture

Following isolation of choroids (described above), choroids from 15 day old chicks were transferred to sterile 48-well plates containing 300 μl culture medium [1:1 mixture of Dulbecco’s Modified Eagle’s Medium and Ham’s F12 containing streptomycin (0.1 mg/ml), penicillin (100 units/ml) gentamicin (50 μg/ml)] in the presence of the NO donor, PAPA-NONOate (1.5 mM in culture medium; Cayman Chemical, Ann Arbor, MI), or culture medium alone in a humidified incubator with 5% CO_2_, overnight at 37 °C. Following incubation, choroids were snap frozen and RNA isolated for RNA-seq and TaqMan real time PCR assays.

### RNAseq

Choroids were isolated from the left and right eyes of 5 chicks as described above and placed in 48-well plates containing 300 μl culture medium [1:1 mixture of Dulbecco’s Modified Eagle’s Medium and Ham’s F12 containing streptomycin (0.1 mg/ml), penicillin (100 units/ml) and gentamicin (50 μg/ml)] in the presence of the NO donor, PAPA-NONOate (1.5 mM in culture medium; Cayman Chemical, Ann Arbor, MI), or culture medium alone in a humidified incubator with 5% CO_2_, overnight at 37 °C. Following incubation, choroids were snap frozen in liquid nitrogen and stored at −80°C. Total RNA was isolated using TRIzol reagent (ThermoFisher Scientific) followed by DNase treatment (DNA-free, Applied Biosystems) as described previously [30]. RNA concentration and purity were determined via the optical density ratio of 260/280 using a Nanodrop ND-1000 spectrophotometer and stored at −80 °C until use.

### Illumina RNAseq libraries

Stranded RNA-seq libraries were constructed using NEB Next poly(A) mRNA isolation kit with the SWIFT RNA Library Kit and the established protocols. The library construction was done using 100 ng of RNA. Each of the libraries was indexed during library construction in order to multiplex for sequencing. Samples were normalized and the 10 libraries were pooled onto a 150 paired end run on Illumina’s NovaSeq Platform. ∼ 56,000,000 – 81,000,000 raw read pairs were collected for each sample. Raw data were analyzed by applying a custom computational pipeline consisting of the open-source gSNAP, Cufflinks, and R for sequence alignment and ascertainment of differential gene expression [31]. Pairwise comparison of the expression results were performed using the total mapping results for control vs treated samples using a paired T-test to control for individual variance. Differential gene lists were created with ≥ 2 fold expression cutoff and significant p-values of < 0.05 were submitted to pathway analysis using Ingenuity Pathway Analysis (Qiagen, Germantown, MD) to identify pathways of interest that were modified by treatment with PAPA-NONOate.

### TaqMan Quantitative PCR (RT-Quantitative PCR)

Choroids were isolated from the left and right eyes of 9 chicks as described above and placed in 48-well plates containing 300 μl culture medium [1:1 mixture of Dulbecco’s Modified Eagle’s Medium and Ham’s F12 containing streptomycin (0.1 mg/ml), penicillin (100 units/ml) and gentamicin (50 μg/ml)] in the presence of the NO donor, PAPA-NONOate (1.5 mM in culture medium; Cayman Chemical, Ann Arbor, MI), or culture medium alone in a humidified incubator with 5% CO_2_, overnight at 37 °C. Following incubation, choroids were snap frozen in liquid nitrogen and stored at −80°C. Total RNA was isolated using TRIzol reagent (ThermoFisher Scientific) followed by DNase treatment (DNA-free, Applied Biosystems) as described previously [30]. RNA concentration and purity were determined via the optical density ratio of 260/280 using a Nanodrop ND-1000 spectrophotometer and stored at −80 °C until use. cDNA was generated from DNase-treated RNA using a High Capacity RNA to cDNA kit. Real time PCR was carried out using a Bio-Rad CFX 96. 20-μl reactions were set up containing 10 μl of TaqMan 2× Universal Master Mix (Applied Biosystems), 1 μl 20× 6-carboxyfluorescein (FAM)-labeled Assay Mix (Applied Biosystems), and 9 μl of cDNA. Each sample was set up in duplicate with specific primers and probed for chicken cell division cycle associated 3 (CDCA3, assay ID number Gg07161292_g1), chicken endothelial cell specific molecule 1 (ESM1, assay ID number APMF4DV), chicken heat shock protein family b (small) member 9 (HSPB9, assay ID number Gg03812884_s1), chicken leucine rich single-pass membrane protein 1 (LSMEM1, assay ID number Gg07167399_m1), chicken solute carrier family 7 member 14 (SLC7A14, assay ID number Gg07192083_m1), chicken STEAP4 metalloreductase (STEAP4, assay ID number Gg07188578_m1) and the reference gene chicken GAPDH (assay ID number Gg03346982_m1) (Thermofisher Scientific). The PCR cycle parameters were an initial denaturing step at 95 °C for 10 min followed by 45 cycles of 95 °C for 15 s and 60 °C for 1 min. Normalized gene expression was determined by the ΔΔ*c*(*t*) method [32] using Bio-Rad CFX Manager^TM^ version 3.1 and reported values represent the average of duplicate samples.

### Statistics

Sample sizes were calculated using G*Power 3.1.9.2 using two tailed tests with an α = 0.05, and an effect size determined by group means and standard deviations previously published by this lab and others [16, 27]. All experiments were repeated at least one time, and sample sizes and results reported reflect the cumulative data for all trials of each experiment. All data was subjected to the D’Agostino & Pearson test to test the normality of the data. Data that passed the D’Agostino & Pearson test were subjected to parametric analyses. Parametric analyses between two groups were made using paired or unpaired Student’s t-tests. Data that failed the D’Agostino & Pearson test, or had a sample size too small for the D’Agostino & Pearson normality test were subjected to non-parametric analyses. Non-parametric tests between two groups were made using the Wilcoxon signed rank test for matched pairs (GraphPad Prism 5, La Jolla, CA). Results were considered significant with p-value ≤ 0.05.

## RESULTS

### NO Imaging in the Choroid

The NO-sensitive dye DAF-2DA is known to be a very sensitive and specific indicator for NO production in the retina [28,33]. We therefore used this dye to image NO-containing cells in choroids of chick eyes under a visual condition (recovery from induced myopia) previously shown to be dependent on NO [24]. In recovering choroids, numerous DAF-2DA positive cells were visualized within choroid whole mounts (**Figures 1C, 1D, 1H and 1I**). DAF-2DA positive cells were variable in shape; some with long processes extending from one or both ends of the cell bodies. These cells were concentrated on the proximal side of the choroid, near the RPE, and appeared to be distributed throughout the choroidal stroma. Very few labelled cells were detected in contralateral control choroids (**Figure 1A, 1B, 1F and 1G**). No green fluorescence was observed in tissue that was not treated with DAF-2DA (**Figures 1E and 1J**). These results suggest increased intracellular concentrations of NO in a subpopulation of choroidal cells during recovery from induced myopia.

**Figure 1.**
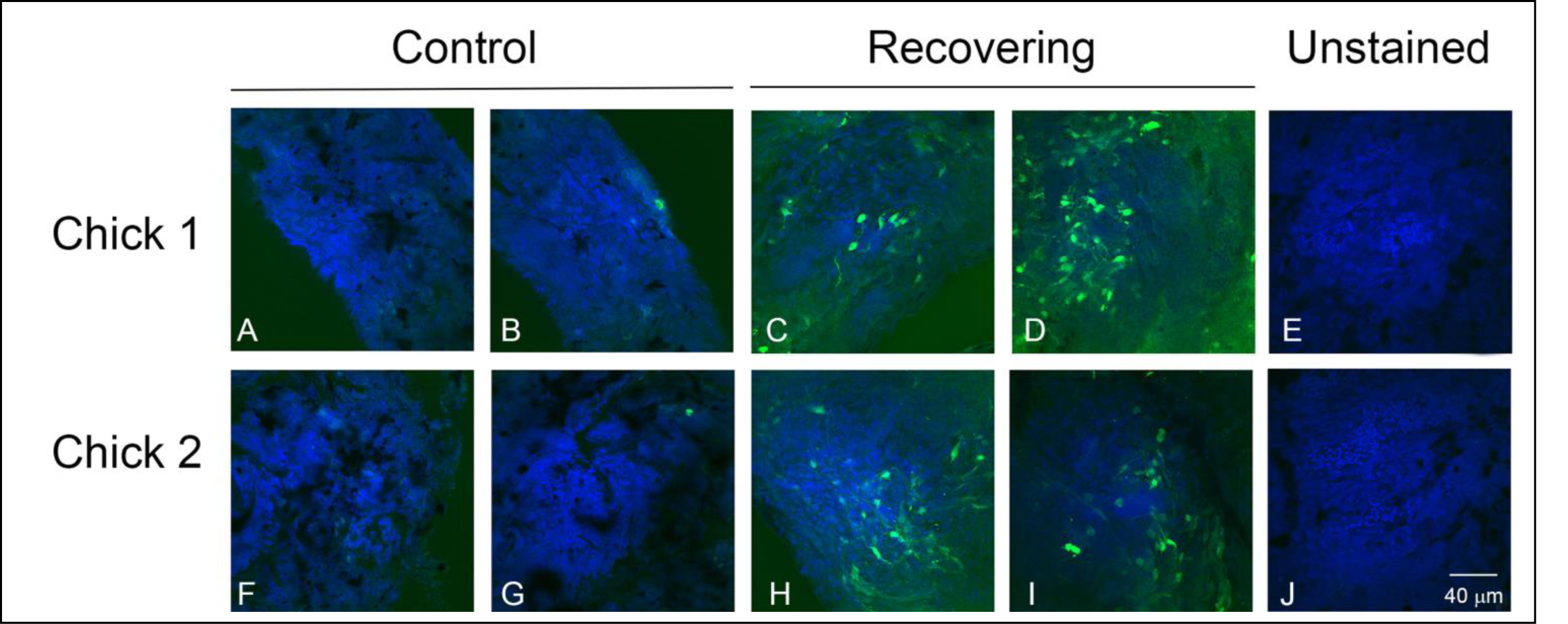
Imaging of intracellular NO with DAF-2DA in chick choroid whole mounts from chick eyes following 3 hrs of recovery from induced myopia and from contralateral control eyes. DAF-2DA labelled cells (green) are readily identified in recovering choroids, but are largely absent in contralateral control choroids. Nucleii are labelled with DAPI. Bar = 40 μM for all images.

### NO induces significant changes in choroidal gene expression

Treatment of normal chick choroids with the NO donor, PAPA-NONOate, resulted a total of 837 differentially expressed genes compared with untreated choroids from the contralateral eyes (fold change ≥ 1.09 and ≤ −1.07 P < 0.05, paired t-test). 259 genes were found to be upregulated and 578 genes were downregulated (**Supplementary Table S1**). A heat map and volcano plot representing all differentially expressed genes are shown in **Figure 2**.

**Figure 2.**
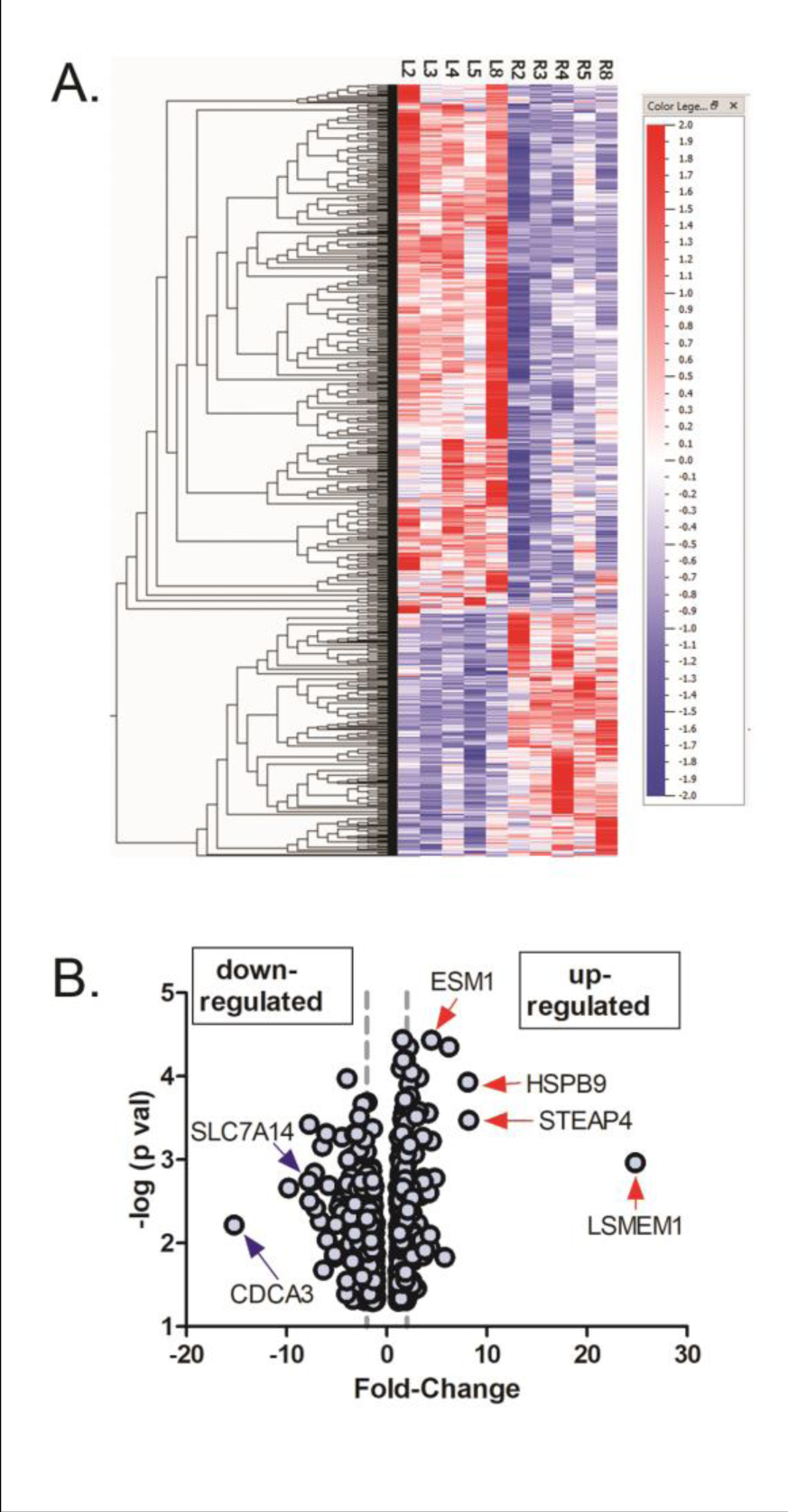
Differentially expressed genes in chick choroids in response to 1.5 mM PAPA-NONOate. **(A)** Heat map showing signal intensities of differentially expressed genes in control (L2, L3, L4, L5, and L8) and PAPA-NONOate-treated (R2, R3, R4, R5, and R8) choroids. **(B)** Volcano plot of differentially expressed genes in response to PAPA-NONOate treatment (p < 0.05, paired t-test). The x-axis represents the fold change values, and the y-axis represents the negative log of the significance level (paired t-test). The left and right dashed vertical lines indicate positions for 2 fold down regulated genes and 2 fold up regulated genes, respectively. Genes indicated with blue arrows (down-regulated) and red arrows (up-regulated) were confirmed with reverse transcription qPCR.

The top five upregulated genes were LSMEM1, STEAP4, HSPB9, HSPA2, and CCL19 (**Table 1**), whereas the top five downregulated genes were CDCA3, SMC2, a novel gene (ENSALGALG00000050836), an uncharacterized gene (LOC107054158), and SPAG5 (**Table 2**).

**Table 1.** 10 Most Overexpressed Genes in the RNA-seq transcriptome of chick choroids treated with 1.5 mM PAPA-NONOate.

|  |  |  |  |  |  |
| --- | --- | --- | --- | --- | --- |
| **Accession number** | **Ensembl ID** | **FPKM control (L) ± SEM** | **FPKM treated (R) ± SEM** | **Fold Change** | **Gene ID** |
| XM040659395 | ENSGALG00000009446 | 0.24432 ± 0.071700814 | 5.96130 ± 1.699807346 | 24.83 | LSMEM1 |
| XM040665708 | ENSGALG00000008997 | 3.02881 ± 0.71931704 | 24.73888 ± 3.790289 | 8.17 | STEAP4 |
| XM425870 | ENSGALG00000023818 | 101.19700 ± 14.0261479 | 820.09240 ± 166.6266 | 8.1 | HSPB9 |
| XM421406 | ENSGALG00000011715 | 96.47044 ± 10.3197415 | 597.89660 ± 56.0171 | 6.2 | HSPA2 |
| XM424980 | ENSGALG00000046192 | 3.11683 ± 0.53857 | 18.0959 ± 5.35255 | 5.8 | CCL19 |
| XM419084 | ENSGALG00000013575 | 1.94991 ± 0.46951024 | 9.36739 ± 1.58115 | 4.81 | IFI6 |
| XM428357 | ENSGALG00000027707 | 1.58904 ± 0.14123103 | 7.07570 ± 0.019873 | 4.45 | ESM1 |
| XM424455 | ENSGALG00000043734 | 4.49535 ± 0.58149042 | 20.02234 ± 1.858336 | 4.45 | LIPG |
| XM003642832 | ENSGALG00000023819 | 9.27436 ± 2.72539 | 40.4566 ± 6.04158 | 4.36 | HSP30C (predict |
| XM421662 | ENSGALG00000045085 | 33.2954 ± 6.4383 | 143.388 ± 32.3406 | 4.31 | IFIT5 |

**Table 2.** 10 Most Underexpressed Genes in the RNA-seq transcriptome of chick choroids treated with 1.5 mM PAPA-NONOate.

|  |  |  |  |  |  |
| --- | --- | --- | --- | --- | --- |
| **Accession number** | **Ensembl ID** | **FPKM control (L) ± SEM** | **FPKM treated (R) ± SEM** | **Fold Change** | **Gene ID** |
| XM001232833 | ENSGALG00000014513 | 13.56922 ± 2.87044982 | 0.8922052 ± 0.267956974 | -15.25 | CDCA3 |
| XM040655444 | ENSGALG00000015691 | 21.53703 ± 3.37073041 | 2.20113 ± 14.25296 | -9.79 | SMC2 |
| - | ENSGALG00000050836 | 8.73492 ± 2.3000145 | 1.11604 ± 6.2052 | -7.79 | Novel gene |
| XM040706642 | ENSGALG00000055004 | 18.35300 ± 4.12570678 | 2.37274 ± 13.43117 | -7.74 | Uncharacterized I<br>(predicted) |
| XM040687237 | ENSGALG00000048771 | 8.415996 ± 1.035199107 | 1.0873174 ± 0.252546471 | -7.72 | SPAG5 (predicted |
| XM040679569 | ENSGALG00000009349 | 7.45987 ± 1.17821557 | 1.00821 ± 2.384324 | -7.39 | SLC7A14 |
| XM040668831 | ENSGALG00000016442 | 23.06680 ± 3.3598115 | 3.20059 ± 14.08091 | -7.21 | RRM2 |
| XM040689493 | ENSGALG00000005961 | 25.57842 ± 4.826339933 | 3.601136 ± 0.440039301 | -7.11 | DGUOK |
| XM015288064 | ENSGALG00000003085 | 28.17486 ± 4.72625028 | 4.229584 ± 0.840515731 | -6.66 | CDK1 |
| NM001030731 | ENSGALG00000008439 | 14.71402 ± 1.520979489 | 2.289716 ± 0.233476644 | -6.42 | CD36 |

### Enriched canonical pathways detected using IPA

The complete list of significantly enriched canonical pathways is included in **Supplementary Table S2**. A total of 78 enriched canonical pathways were identified by applying the −log (P-value) >2 threshold, and the top 25 of the 78 representative pathways ranked according to their −log (P-value) (**Figure 3**).

**Figure 3.**
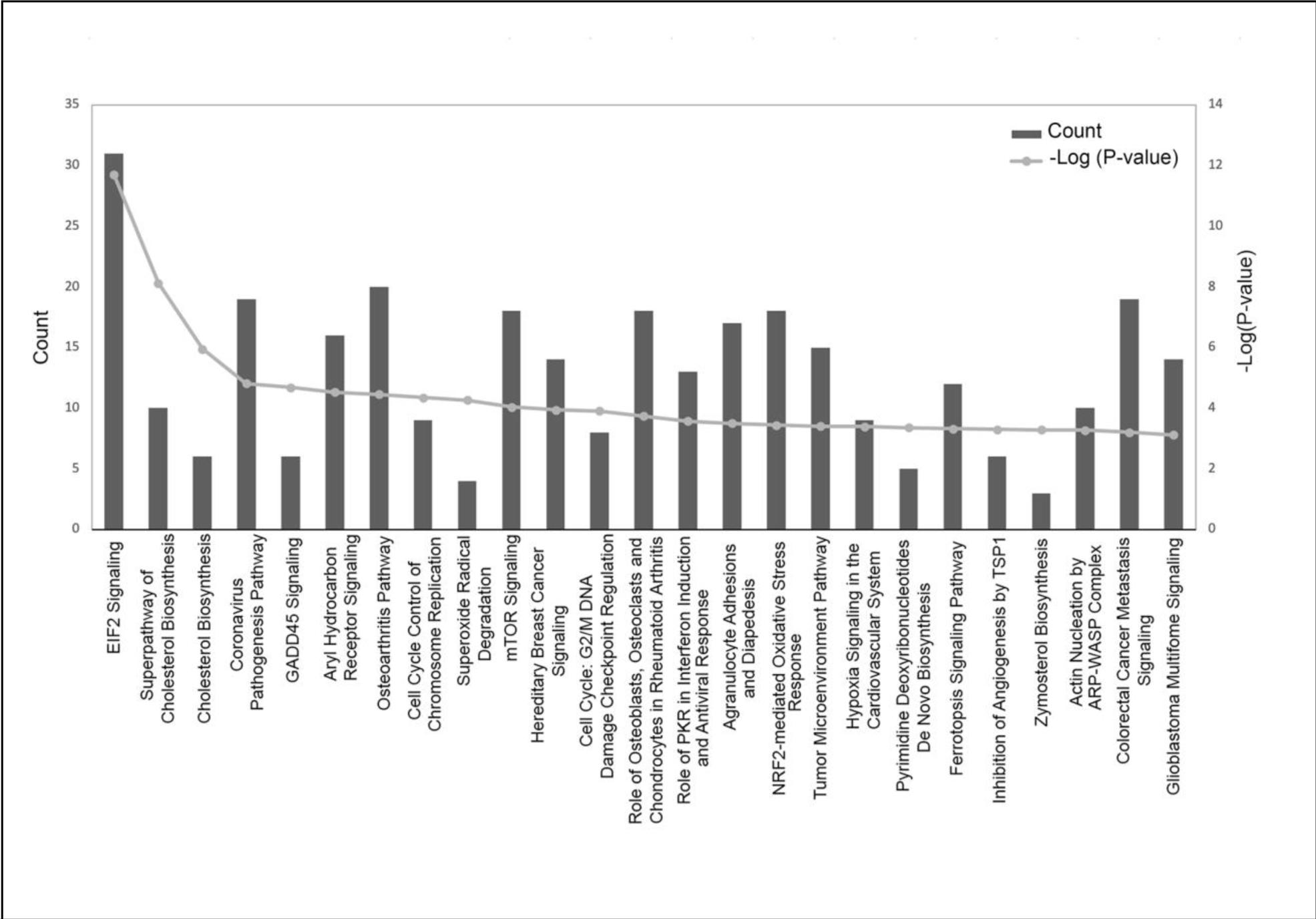
Histogram of major canonical pathways. The left y-axis (“Count”) represents the number of genes counted for each of the listed canonical pathways and the right y-axis represents the −log (P-value). Pathways are sorted left to right by descending P-values. EIF2, eukaryotic initiation factor 2; GADD45, growth arrest and DNA damage-inducible 45; mTOR, mammalian target of rapamycin; PKR, protein kinase RNA-activated; NRF-2, nuclear factor erythroid 2–related factor 2; TSP1, thrombospondin 1; ARP-WASP, actin-related proteins - Wiskott–Aldrich syndrome protein.

The “E1F2 Signaling” pathway was the highest-ranking signaling pathway with a −log(P-value) of 11.703. Other significantly enriched pathways included “Superpathway of Cholesterol Biosynthesis” [-log(P-value) of 8.12], “Cholesterol Biosynthesis” [-log (P-value) of 5.955], and “Coronovirus Pathogenesis Pathway” [-log (P-value) of 4.812].

### Classification of diseases and functions mediated by NO

The role of NO in various diseases and in cellular function was next determined based on the −log (P-values). A complete list of disease and function classifications are provided in **Supplementary Table S3**. The top 28 categories with their −log(P-value) and number of associated genes in our dataset (count) are shown in **Figure 4A** sorted by z score (a numerical indicator of how closely the observed changes in gene expression match the predicted up/down gene regulation patterns for each pathway, disease or function, based on the literature). “Cell Death and Survival” had the highest z-score range (1.608 – 3.470) whereas “Cell Movement” had the lowest z-score range of (−3.221 – 1.083). The major disease categories with the highest −log (P-values) and their associated pathophysiological processes are shown in **Figures 4B - D**. A representative heatmap of the major diseases and functions with all underlying pathophysiological processes is shown in **Supplemental Figure S1**. NO was found to alter choroidal gene expression patterns similar to that of a number of diseases and cellular functions, including “Organismal Injury and Abnormalities (Z-score range of −3.121 – 3.3) (**Figure 4B**), “Cell Death and Survival” (Z-score range of −1.608 −3.47) (**Figure 4C**), and “Cell Movement” (Z-score range = −3.221 – 1.083) (**Figure 4D**). Additional cellular functions affected by NO included “Cellular Development” (Z-score range=-1.555 – 1.844), “Tissue Development” (Z-score range = −1.555 – 1.844) and “Cellular Growth and Proliferation” (Z-score = −1555 – 1.611).

**Figure 4.**
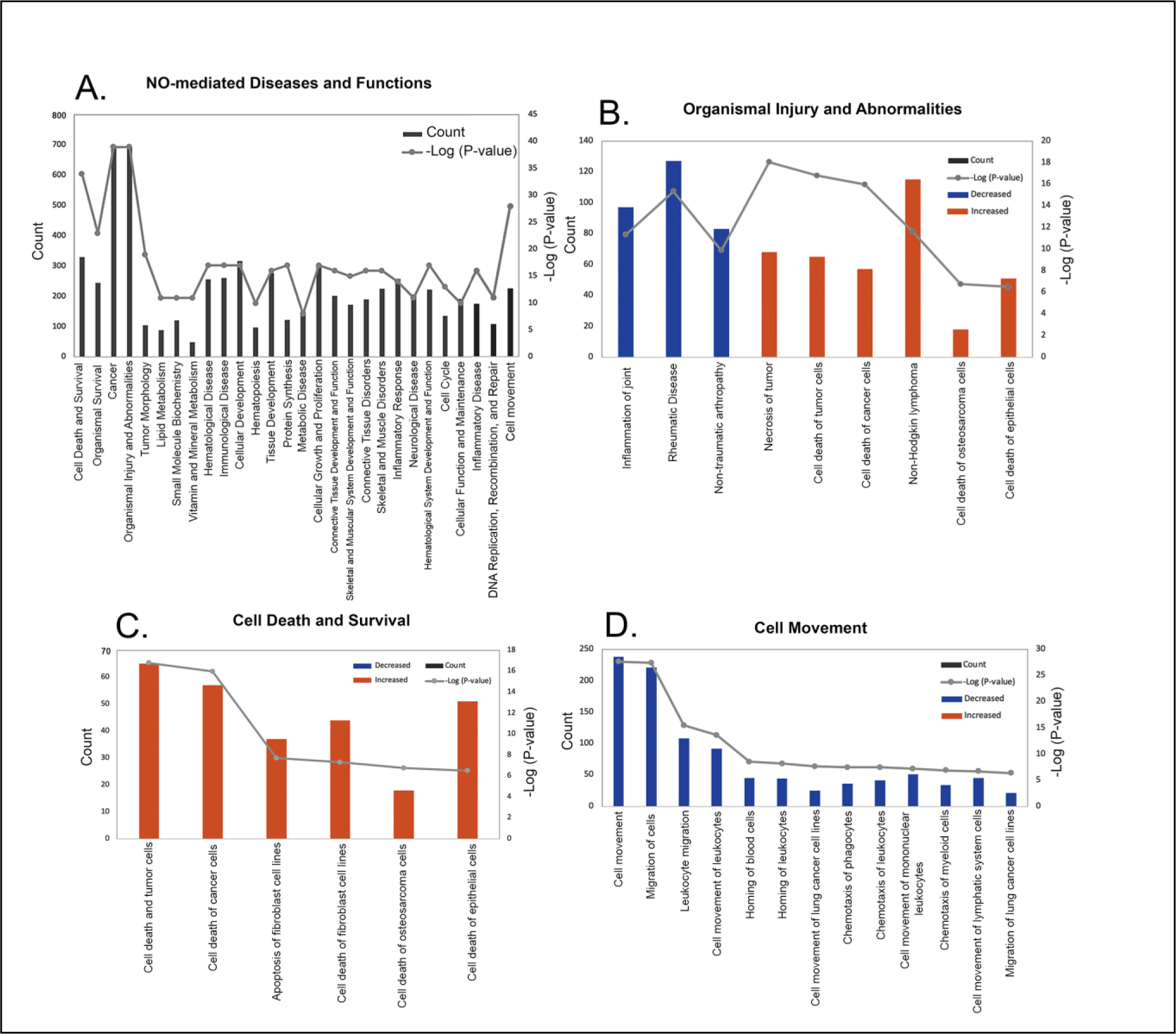
Classification of diseases and functions mediated by nitric oxide (NO) in the choroid. Categories are shown in term of the −log (P-value; right y-axis) and the number of differentially expressed genes counted (left y-axis). **(A)** 28 representative classifications of diseases and functions mediated by NO. **(B)** Classification of pathophysiological processes mediated by NO related to organismal injury and abnormalities. **(C)** Classification of pathophysiological processes mediated by NO related to cell death and survival. **(D)** Classification of possible functions of NO in the choroid related to cell movement.

### Regulatory effects analysis

The regulatory effects analysis demonstrates the possible pathways of upstream regulatory networks and downstream functions that involve the gene expression changes induced by NO (**Figure 5**). In total, 60 types of regulatory effects were found (**Supplementary Table S4**), ranked by their Consistency Scores. The Consistency Score is an indicator used to describe the causal consistency of the upstream regulatory factor in the network, the dataset and dense connection metric between disease and function [34]. Among them, the highest ranked regulatory effect with a consistency score of 16.364 is shown in **Figure 5**. The Regulatory effects analysis strongly suggests that NO induces or is predicted to induce gene expression changes in SQSTM1, CXCL8, ILK, ST8SIA1, CASP3, NEDD4, GADD45A and SPDEF, that may be involved in the chemotaxis of myeloid cells, chemotaxis of phagoctyes, migration of lung cancer cell lines, cell movement of mononuclear leukocytes, and cell movement of lymphatic system cells mainly through mediating their targets, including CDKN1A, MMP7, VIM, TGFB1, IL1B, CD74, LGALS3, HMOX1, NFKBIA, MMP9, NQO1, ACKR3 and ITGA3, whose gene expression was shown to be altered by NO in our study (**Figure 5**).

**Figure 5.**
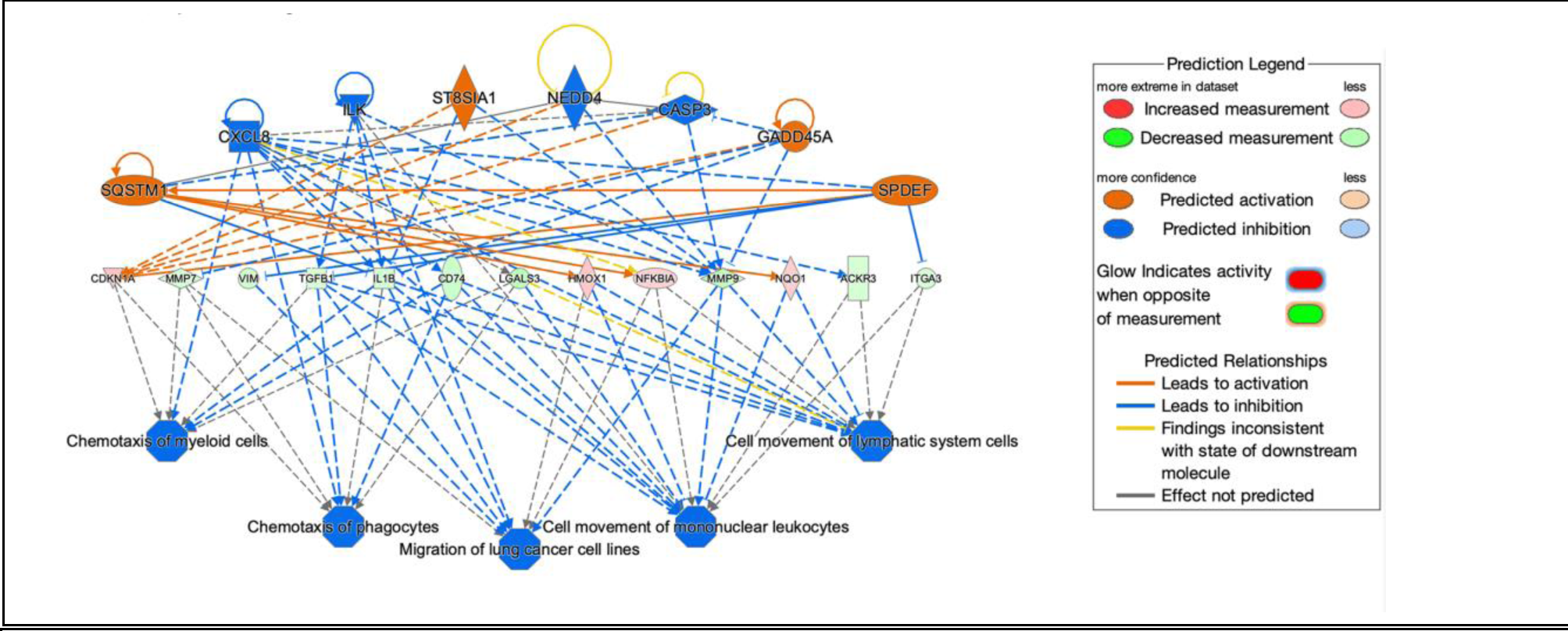
Network diagram representing the highest ranked regulatory effect pathway of PAPA-NONOate on choroidal gene expression based on the consistency score. Upregulated genes are labeled in red or pink, while downregulated genes are labeled in green. The saturation of color is correlated with the fold change of the gene (high saturation means high fold change and low saturation means low fold change). Solid lines indicate direct connections, while dotted lines indicate indirect connections (circular arrows indicate autocrine influence). Triangles represent kinases, horizontal diamonds represent peptidases, dashed squares represent growth factors, squares represent cytokines, vertical ovals represent transmembrane receptors, horizontal ovals represent transcription regulators, vertical diamonds represent enzymes, rectangles represent G-protein coupled receptors, and circles represent “other”. Treatment of choroids with PAPA-NONOate is predicted to inhibit pathways involved in chemotaxis of myeloid cells and phagocytes, migration of lung cancer cell lines, and cell movement of mononuclear leukocytes and lymphatic system cells.

### Validation of target genes

Six genes expressed in the choroid that demonstrated the highest fold changes (up and down regulated) (indicated in **Figure 2B**) were selected for qPCR validation. The genes CDCA3 and SLC7A14 where shown to be significantly downregulated both by RNAseq and Taqman™ real time PCR, while genes ESM1, HSPB9, LSMEM1, and STEAP4 were significantly upregulated by treatment with PAPA-NONOate (**Figure 6**).

**Figure 6.**
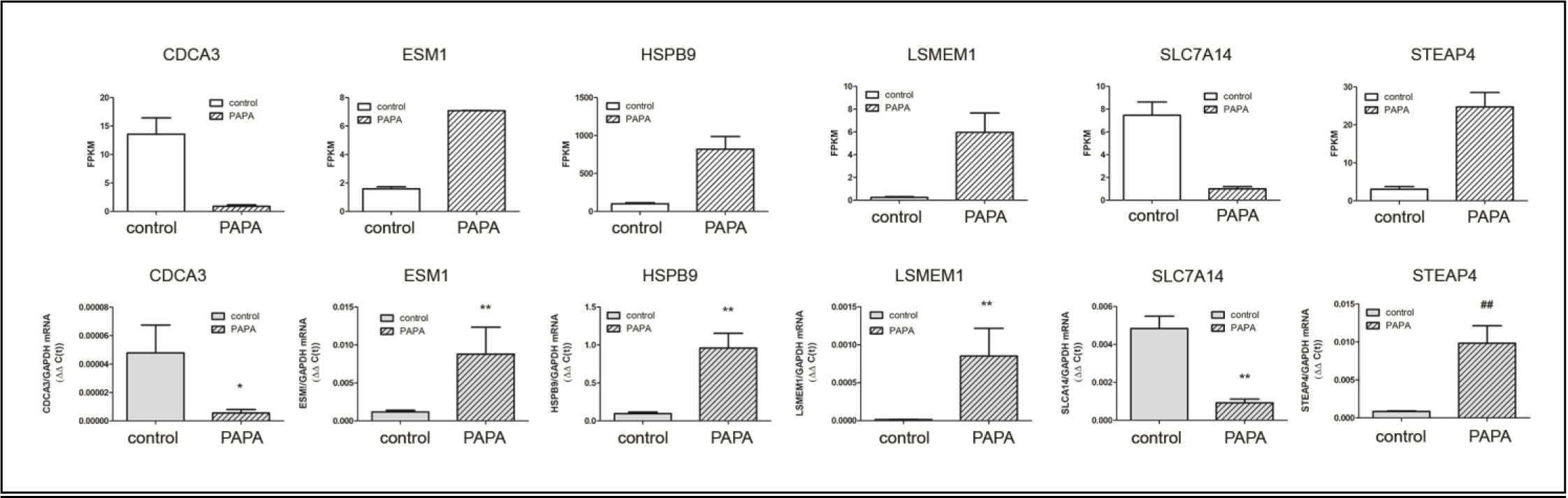
Confirmation of RNA-Seq results by qRT-PCR analysis. A. mRNA expression of six target genes, identified as significantly up- or down regulated by NO by RNA-seq, was validated using Taqman reverse transcription-quantitative PCR. A) RNA seq results, expressed as fragments per kilobase of transcript per million mapped reads [(FPKM); p< 0.05 for all genes, paired t-test]. B) Taqman RT-qPCR results, expressed as the gene of interest, normalized to GAPDH (ΔΔC(t)). *p < 0.05; Wilcoxan matched-pairs signed rank test, **p < 0.01; Wilcoxan matched-pairs signed rank test, ##p < 0.01; Student’s t-test for matched pairs.

## DISCUSSION

NO is now recognized as a major signaling molecule in eukaryotes that controls diverse cellular processes, including immune responses, angiogenesis, metastasis, and apoptosis [35–38]. Under normoxic or hyperoxic conditions, nitric oxide is endogenously synthesized in various tissues by the transformation of L-arginine to L-citrulline by three distinct isoforms of nitric oxide synthase (NOS). The neuronal (nNOS) and endothelial (eNOS) isoforms are calcium-dependent isoforms that yield low fluxes of NO in the pico-to nanomolar range for a short period of time varying from seconds to minutes for regulating intercellular and extracellular signaling pathways [39]. Inducible nitric oxide synthase (iNOS) is the calcium-independent isoform and is induced by cytokines, endotoxins, and hypoxia under oxidative stress. iNOS produces significant concentrations (micromolar range) of NO for a longer period of time ranging from hours to days [40, 41].

Several studies suggest that NO may be involved in the regulation of postnatal eye growth. Topical or intravitreal administration of the NOS substrate, L-arginine, or the NO donor, SNP, inhibits ocular elongation and myopia development [25, 42], while administration of the nonspecific NOS inhibitors, L-NAME and NG-monomethyl-L-arginine acetate (L-NMMA) as well as the nNOS-specific inhibitor, N(omega)-propyl-L-arginine (NPA), block the recovery from induced myopia [23, 24, 43]. Additionally, the NOS inhibitors, L-NIO, L-NMMA, and NPA have been shown to block the myopia-inhibiting effects of muscarinic receptor antagonist, atropine, and the dopamine agonist, quinpirole, suggesting that the effects these myopia-inhibiting compounds are mediated by NO [25, 44]. Therefore, it is of great interest to elucidate the role of NO in ocular growth control in order to gain insight into the fundamental mechanisms underlying visually guided ocular growth.

Many cell types in the eye are capable of synthesizing NO. On the ocular surface, NO is synthesized by corneal epithelial cells, fibroblasts, endothelium and inflammatory cells [45, 46]. nNOS-like immunoreactivity can be found in all major cell types in the chick retina and in intrinsic choroidal neurons [47–49]. Additionally, eNOS is associated with choroidal endothelium [50] and iNOS has been shown to be synthesized by the RPE [51], and by choroidal macrophages [52]. Our NO imaging using the intracellular NO indicator, DAF-2DA, indicates that a population of extravascular choroidal cells, located in the proximal choroid, contain NO. It is unclear as to whether these DAF-2DA - positive cells are the site of choroidal NO synthesis, or if they represent the targets of NO via paracrine signaling from another cell type.

In the present study, the NO-donor, PAPA-NONOate was used to study the downstream effects of NO on choroidal gene expression. PAPA NONOate spontaneously dissociates in a pH-dependent, first-order process with a half-life of 15 minutes at 37°C, respectively, (pH 7.4) to liberate 2 moles of NO per mole of parent compound [53]. Treatment of choroids with PAPA-NONOate resulted in the upregulation of 259 genes and downregulation of 578 genes. These NO-induced changes in choroidal gene expression were analyzed using the Ingenuity Pathway Analysis™ (IPA) application to uncover the signaling pathways, interactions and functional roles of differentially expressed eyes associated with NO treatment. The results of the transcriptome analysis are summarized in the form of a model showing the major functional pathways and the differentially expressed genes that were either activated or inhibited in the choroid after PAPA-NONOate treatment (**Figure 7**).

**Figure 7.**
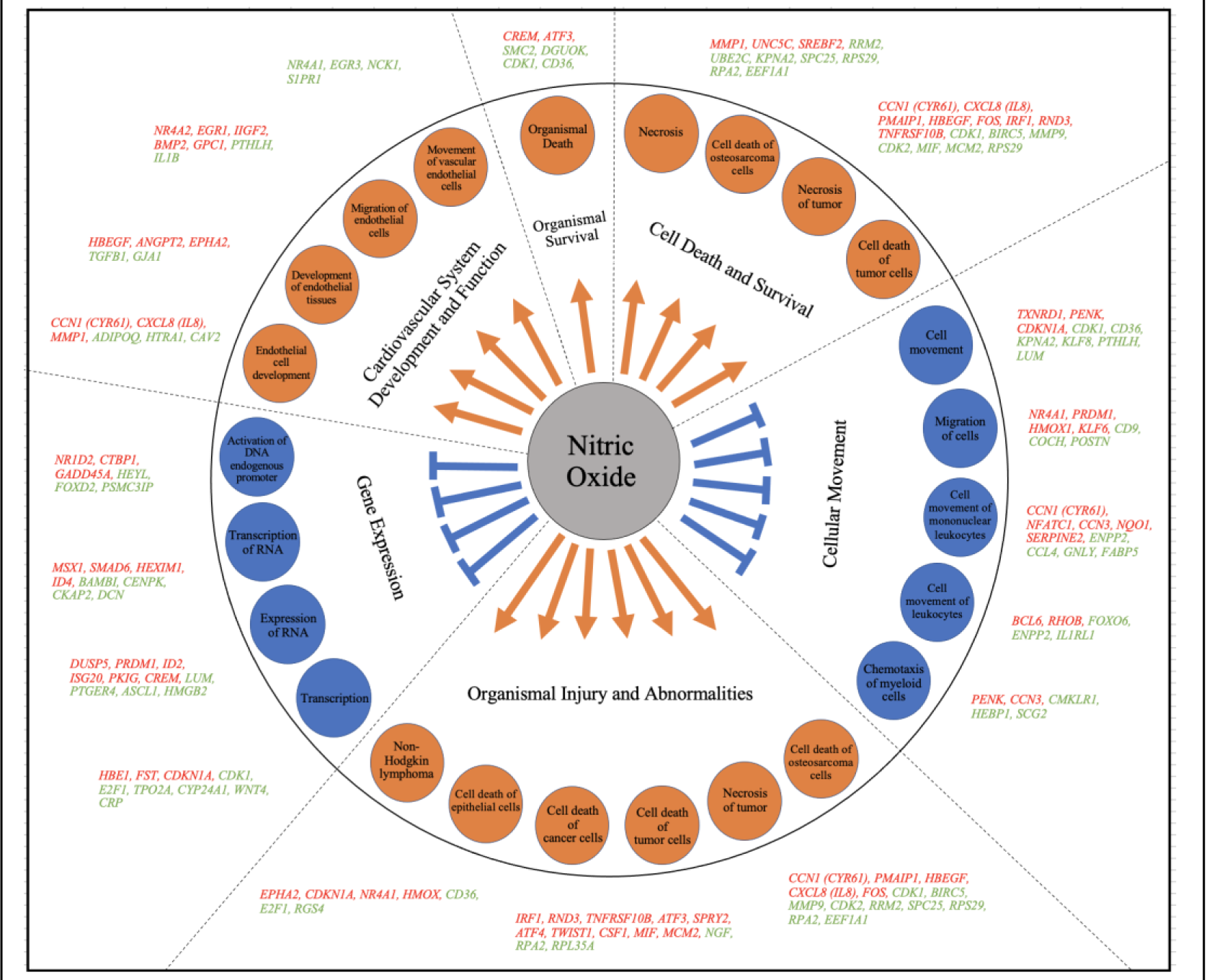
Model summarizing the different regulatory pathways and their respective differentially expressed genes in chick choroid following treatment with PAPA-NONOate. Ingenuity analysis identified differentially expressed genes (up-regulated; orange font, down-regulated; green font) involved in important physiological activities such as; organismal survival, cell death and survival, cellular movement, organismal injury and abnormalities, gene expression and cardiovascular system development and function. Within each of the above categories, several representative sub-categories were identified (in orange and blue circles).

Canonical pathway analysis identified “E1F2 Signaling” as the highest-ranking signaling pathway in the choroid affected by NO. NO has been shown to activate multiple serine-threonine kinases that phosphorylate and activate the alpha subunit of eukaryotic initiation factor 2 (eIF2α) to coordinately regulate physiological and pathological events such as the innate immune response and cell apoptosis [54]. NO-induced activation of eIF2α phosphorylation has been shown to inhibit protein synthesis and cause cell cycle arrest in nonerythroid cells as well as lead to ER-stress, protein degradation and apoptosis. Additionally, NO-induced activation of the eIF2α kinase, EIF2AK3, regulates Ca^2+^ flux and therefore affects cardiac and skeletal muscle contraction.

The NO-induced changes in choroidal gene expression were similar to gene expression changes associated with several diseases and cellular functions. “Organismal Injury and Abnormalities”, “Cell Death and Survival”, and “Cell Movement” were three categories with the lowest p-values (most significantly affected), with NO generally causing activation of cell death, apoptosis, and necrosis, and inhibition of cell movement, migration and inflammation. Based on studies using cancer cell lines, low concentrations of NO donors promote cancer progression by the activation of several mitogenic pathways, including extracellular signal-regulated kinase mTOR and Wnt/Catenin pathways [55–57] whereas higher doses of NO donors (> 600 nM) negatively regulate the proliferation of cancer cells by attenuating the expression of oncogenes and by the upregulation of tumor suppressors (BRCA1/Chk1/p53) culminating in cell cycle arrest via the activation of cell cycle checkpoints [58]. Based on the observed gene expression changes and the concentration of PAPA-NONOate used in the present study (1.5 mM) we predict the concentration of NO in our choroidal organ cultures was on the high end of the NO flux scale (although not directly measured in the present study).

A total of 60 regulatory effects of NO on choroids were predicted using the IPA application. The top regulatory network (based on its consistency score of 16.364) predicted inhibition of the pathways: “chemotaxis of myeloid cells”, “chemotaxis of phagocytes”, “migration of lung cancer cell lines”, “cell movement of mononuclear leukocytes”, and “cell movement of lymphatic system cells”. The choroidal gene expression changes in our dataset driving these networks included increased expression of the genes SQSTM1, CXCL8 (IL8), GADD45A, CDKN1A, HMOX1, NFKBIA and NQO1 and decreased expression of MMP7, VIM, TGFB1, IL1B, CD74, LGALS3, MMP9, ACKR3, and ITGA3.

Six genes were selected for Taqman™ qPCR validation which were either highly upregulated or highly downregulated. These genes have been shown to be involved in a variety of cell functions including the regulation of cell cycle, cell proliferation, protein ubiquitination, amino acid transport and copper ion transport. The genes, ESM1, HSPB9, LSMEM1, and STEAP4 were confirmed to be significantly upregulated, while genes CDCA3 and SLC7A14 were significantly downregulated. Not surprisingly, several of these genes have previously been shown to be involved in NO synthesis, cytokine signaling and inflammation. Among these, the upregulated gene, endothelial cell-specific molecule 1 (ESM1, endocan) encodes a secreted protein that has been shown to be expressed in endothelial cells in the human lung and kidney tissues. The expression of this gene is regulated by cytokines [59, 60], and is known to induce vascular inflammation [61, 62]. Additionally, ESM1 has been shown to promote nitric oxide production through iNOS activation [63].

The gene, six transmembrane epithelial antigen of prostate (STEAP) 4, has been shown to be upregulated by TNF-α and other inflammatory cytokines such as IL-6 and IL-1β [64]. The STEAP4 protein has been shown to maintain iron and copper homeostasis by reducing Fe^3+^ to Fe^2+^ and Cu^2+^ to Cu^+^ and promoting their transmembrane transport [65]. Furthermore, some studies have shown that STEAP4 plays a protective role against inflammatory damage in hepatocytes [66], mesangial cells [67], and synovial cells [68].

The downregulated gene, solute carrier family 7, member 14 (SLC7A14) is a cationic amino acid transporter. In some cells, SLC7A14 may regulate the rate of NO synthesis by controlling the uptake of l-arginine as the substrate for NOS. Mutations in the *SLC7A14* gene cause autosomal recessive retinitis pigmentosa (arRP) [69]. *Slc7a14* has been shown to be highly expressed in neuronal tissues, including the brain, spinal cord and retina, and that the expression levels increased during early embryogenesis [70]. Interestingly, Knockdown of *Slc7a14* led to dose-dependent microphthalmia that was reversed by overexpression, suggesting a role in ocular growth [69].

### Conclusions

We investigated choroidal gene expression changes in response to treatment with the NO donor PAPA-NONOate, in order to gain insights into the role of NO in the choroidal response associated with visually induced changes in eye growth. The findings reported herein indicate that NO induced changes in choroidal gene expression that would be predicted to inhibit pathways involved in cell proliferation, cell movement, and gene expression. In contrast, NO treatment was predicted to activate pathways involved in cell and organismal death, necrosis, and cardiovascular system development. Considering that NO signaling is a potential target for the treatment of myopia and other ocular diseases, the results of this study may provide insights into possible choroidal effects of therapies designed to increase ocular concentrations of NO.

## Acknowledgements

This work was supported by NIH grant R01EY09391 (JAS) and by NIGMS COBRE Grant P30GM122744 (Ma, J-X., PI). The authors would like to thank Dr. Norman Hord (Department of Nutritional Sciences, University of Oklahoma Health Science Center) and Dr. William Stell (Department of Cell Biology and Anatomy Department of Cell Biology, University of Calgary - Cumming School of Medicine) for their helpful discussions and suggestions.

## References

1. Cyr AR, Huckaby LV, Shiva SS, Zuckerbraun BS. Nitric Oxide and Endothelial Dysfunction. Crit Care Clin. 2020;36:307–21.

2. Sharma JN, Al-Omran A, Parvathy SS. Role of nitric oxide in inflammatory diseases. Inflammopharmacology. 2007;15:252–9.

3. Erdinest N, London N, Ovadia H, Levinger N. Nitric Oxide Interaction with the Eye. Vision (Basel). 2021;5:29.

4. Toda N, Nakanishi-Toda M. Nitric oxide: ocular blood flow, glaucoma, and diabetic retinopathy. Prog Retin Eye Res. 2007;26:205–38.

5. Chiou GC. Review: effects of nitric oxide on eye diseases and their treatment. J Ocul Pharmacol Ther. 2001;17:189–98.

6. Wildsoet CF. Active emmetropization--evidence for its existence and ramifications for clinical practice. Ophthalmic Physiol Opt. 1997;17:279–90.

7. Troilo D, Smith EL, 3rd, Nickla DL, Ashby R, Tkatchenko AV, Ostrin LA, Gawne TJ, Pardue MT, Summers JA, Kee CS, Schroedl F, Wahl S, Jones L. IMI - Report on Experimental Models of Emmetropization and Myopia. Invest Ophthalmol Vis Sci. 2019;60:M31–M88.

8. Mutti DO. Hereditary and environmental contributions to emmetropization and myopia. Optom Vis Sci. 2010;87:255–9.

9. Organization WH. The impact of myopia and high myopia. WHO. 2016; Geneva, Switzerland.

10. Dolgin E. The myopia boom. Nature. 2015;519:276–8.

11. Howlett MH, McFadden SA. Form-deprivation myopia in the guinea pig (Cavia porcellus). Vision Res. 2006;46:267–83.

12. Shen W, Vijayan M, Sivak JG. Inducing form-deprivation myopia in fish. Invest Ophthalmol Vis Sci. 2005;46:1797–803.

13. Tejedor J, de la Villa P. Refractive changes induced by form deprivation in the mouse eye. Invest Ophthalmol Vis Sci. 2003;44:32–6.

14. Wallman J, Turkel J, Trachtman J. Extreme myopia produced by modest change in early visual experience. Science. 1978;201:1249–51.

15. Sherman SM, Norton TT, Casagrande VA. Myopia in the lid-sutured tree shrew (Tupaia glis). Brain Res. 1977;124:154–7.

16. Wallman J, Adams JI. Developmental aspects of experimental myopia in chicks: susceptibility, recovery and relation to emmetropization. Vision Res. 1987;27:1139–63.

17. Summers JA, Schaeffel F, Marcos S, Wu H, Tkatchenko AV. Functional integration of eye tissues and refractive eye development: Mechanisms and pathways. Exp Eye Res. 2021;209:108693.

18. Wallman J, Wildsoet C, Xu A, Gottlieb MD, Nickla DL, Marran L, et al. Moving the retina: choroidal modulation of refractive state. Vision Res. 1995;35:37–50.

19. Pendrak K, Papastergiou GI, Lin T, Laties AM, Stone RA. Choroidal vascular permeability in visually regulated eye growth. Exp Eye Res. 2000;70:629–37.

20. Rada JA, Palmer L. Choroidal regulation of scleral glycosaminoglycan synthesis during recovery from induced myopia. Invest Ophthalmol Vis Sci. 2007;48:2957–66.

21. Fitzgerald ME, Wildsoet CF, Reiner A. Temporal relationship of choroidal blood flow and thickness changes during recovery from form deprivation myopia in chicks. Exp Eye Res. 2002;74:561–70.

22. Summers JA. The choroid as a sclera growth regulator. Exp Eye Res. 2013;114:120–7.

23. Nickla DL, Wildsoet CF. The effect of the nonspecific nitric oxide synthase inhibitor NG-nitro-L-arginine methyl ester on the choroidal compensatory response to myopic defocus in chickens. Optom Vis Sci. 2004;81:111–8.

24. Nickla DL, Wilken E, Lytle G, Yom S, Mertz J. Inhibiting the transient choroidal thickening response using the nitric oxide synthase inhibitor l-NAME prevents the ameliorative effects of visual experience on ocular growth in two different visual paradigms. Exp Eye Res. 2006;83:456–64.

25. Carr BJ, Stell WK. Nitric Oxide (NO) Mediates the Inhibition of Form-Deprivation Myopia by Atropine in Chicks. Sci Rep. 2016;6:9.

26. Summers JA, Martinez E. Visually induced changes in cytokine production in the chick choroid. Elife. 2021;10.

27. Rada JA, Thoft RA, Hassell JR. Increased aggrecan (cartilage proteoglycan) production in the sclera of myopic chicks. Dev Biol. 1991;147:303–12.

28. Eldred WD, Blute TA. Imaging of nitric oxide in the retina. Vision Res. 2005;45:3469–86.

29. Blute TA, Lee MR, Eldred WD. Direct imaging of NMDA-stimulated nitric oxide production in the retina. Vis Neurosci. 2000;17:557–66.

30. Summers JA, Harper AR, Feasley CL, Van-Der-Wel H, Byrum JN, Hermann M, West CM. Identification of Apolipoprotein A-I as a Retinoic Acid-binding Protein in the Eye. J Biol Chem. 2016;291:18991–9005.

31. Baird NL, Bowlin JL, Cohrs RJ, Gilden D, Jones KL. Comparison of varicella-zoster virus RNA sequences in human neurons and fibroblasts. J Virol. 2014;88:5877–80.

32. Livak KJ, Schmittgen TD. Analysis of relative gene expression data using real-time quantitative PCR and the 2(-Delta Delta C(T)) Method. Methods. 2001;25:402–8.

33. Eldred WD. Real time imaging of the production and movement of nitric oxide in the retina. Prog Brain Res. 2001;131:109–22.

34. Wu C, Chen M, Zhang Q, Yu L, Zhu J, Gao X. Genomic and GeneChip expression profiling reveals the inhibitory effects of Amorphophalli Rhizoma in TNBC cells. J Ethnopharmacol. 2019;235:206–18.

35. Bonavida B, Garban H. Nitric oxide-mediated sensitization of resistant tumor cells to apoptosis by chemo-immunotherapeutics. Redox Biol. 2015;6:486–94.

36. Kielbik M, Szulc-Kielbik I, Nowak M, Sulowska Z, Klink M. Evaluation of nitric oxide donors impact on cisplatin resistance in various ovarian cancer cell lines. Toxicol In Vitro. 2016;36:26–37.

37. Abdel-Messeih PL, Nosseir NM, Bakhe OH. Evaluation of inflammatory cytokines and oxidative stress markers in prostate cancer patients undergoing curative radiotherapy. Cent Eur J Immunol. 2017;42:68–72.

38. Huang Z, Fu J, Zhang Y. Nitric Oxide Donor-Based Cancer Therapy: Advances and Prospects. J Med Chem. 2017;60:7617–35.

39. Somasundaram V, Basudhar D, Bharadwaj G, No JH, Ridnour LA, Cheng RYS, Fujita M, Thomas DD, Anderson SK, McVicar DW, Wink DA. Molecular Mechanisms of Nitric Oxide in Cancer Progression, Signal Transduction, and Metabolism. Antioxid Redox Signal. 2019;30:1124–43.

40. Vannini F, Kashfi K, Nath N. The dual role of iNOS in cancer. Redox Biol. 2015;6:334–43.

41. Kashif K, Duvalsaint, P.L. Nitric Oxide Donors and Therapeutic Applications in Cancer. 1st ed. São Paulo, Brazil: Academic Press; 2017. 75-120 p.

42. McFadden SA. L-Arginine inhibits myopia in the mammalian eye. Invest Ophthalmol Vis Sci. 2021;62:Abs #2278.

43. Nickla DL, Damyanova P, Lytle G. Inhibiting the neuronal isoform of nitric oxide synthase has similar effects on the compensatory choroidal and axial responses to myopic defocus in chicks as does the non-specific inhibitor L-NAME. Exp Eye Res. 2009;88:1092–9.

44. Nickla DL, Lee L, Totonelly K. Nitric oxide synthase inhibitors prevent the growth-inhibiting effects of quinpirole. Optom Vis Sci. 2013;90:1167–75.

45. Kim JC, Cheong TB, Park GS, Park MH, Kwon NS, Yoon HY. The role of nitric oxide in ocular surface diseases. Adv Exp Med Biol. 2002;506(Pt A):687–95.

46. Park JW, Piknova B, Jenkins A, Hellinga D, Parver LM, Schechter AN. Potential roles of nitrate and nitrite in nitric oxide metabolism in the eye. Sci Rep. 2020;10:13166.

47. Wilson M, Nacsa N, Hart NS, Weller C, Vaney DI. Regional distribution of nitrergic neurons in the inner retina of the chicken. Vis Neurosci. 2011;28:205–20.

48. Fischer AJ, Stell WK. Nitric oxide synthase-containing cells in the retina, pigmented epithelium, choroid, and sclera of the chick eye. J Comp Neurol. 1999;405:1–14.

49. Tekmen-Clark M, Gleason E. Nitric oxide production and the expression of two nitric oxide synthases in the avian retina. Vis Neurosci. 2013;30:91–103.

50. Bhutto IA, Baba T, Merges C, McLeod DS, Lutty GA. Low nitric oxide synthases (NOSs) in eyes with age-related macular degeneration (AMD). Exp Eye Res. 2010;90:155–67.

51. Liversidge J, Grabowski P, Ralston S, Benjamin N, Forrester JV. Rat retinal pigment epithelial cells express an inducible form of nitric oxide synthase and produce nitric oxide in response to inflammatory cytokines and activated T cells. Immunology. 1994;83:404–9.

52. Cherepanoff S, McMenamin P, Gillies MC, Kettle E, Sarks SH. Bruch’s membrane and choroidal macrophages in early and advanced age-related macular degeneration. Br J Ophthalmol. 2010;94:918–25.

53. Keefer LK, Nims RW, Davies KM, Wink DA. “NONOates” (1-substituted diazen-1-ium-1,2-diolates) as nitric oxide donors: convenient nitric oxide dosage forms. Methods Enzymol. 1996;268:281–93.

54. Tong L, Heim RA, Wu S. Nitric oxide: a regulator of eukaryotic initiation factor 2 kinases. Free Radic Biol Med. 2011;50:1717–25.

55. Du Q, Zhang X, Liu Q, Zhang X, Bartels CE, Geller DA. Nitric oxide production upregulates Wnt/beta-catenin signaling by inhibiting Dickkopf-1. Cancer Res. 2013;73:6526–37.

56. Garrido P, Shalaby A, Walsh EM, Keane N, Webber M, Keane MM, Sullivan FJ, Kerin MJ, Callagy G, Ryan AE, Glynn SA. Impact of inducible nitric oxide synthase (iNOS) expression on triple negative breast cancer outcome and activation of EGFR and ERK signaling pathways. Oncotarget. 2017;8:80568–88.

57. Lopez-Rivera E, Jayaraman P, Parikh F, Davies MA, Ekmekcioglu S, Izadmehr S, Milton DR, Chipuk, JE, Grimm EA, Estrada Y, Aguirre-Ghiso J, Sikora AG. Inducible nitric oxide synthase drives mTOR pathway activation and proliferation of human melanoma by reversible nitrosylation of TSC2. Cancer Res. 2014;74:1067–78.

58. Van de Wouwer M, Couzinie C, Serrano-Palero M, Gonzalez-Fernandez O, Galmes-Varela C, Menendez-Antoli P, Grau L, Villalobo A. Activation of the BRCA1/Chk1/p53/p21(Cip1/Waf1) pathway by nitric oxide and cell cycle arrest in human neuroblastoma NB69 cells. Nitric Oxide. 2012;26:182–91.

59. Bechard D, Meignin V, Scherpereel A, Oudin S, Kervoaze G, Bertheau P, Janin A, Tonnel A, Lassalle P. Characterization of the secreted form of endothelial-cell-specific molecule 1 by specific monoclonal antibodies. J Vasc Res. 2000;37:417–25.

60. Lassalle P, Molet S, Janin A, Heyden JV, Tavernier J, Fiers W, Devos R, Tonnel AB. ESM-1 is a novel human endothelial cell-specific molecule expressed in lung and regulated by cytokines. J Biol Chem. 1996;271:20458–64.

61. Bechard D, Scherpereel A, Hammad H, Gentina T, Tsicopoulos A, Aumercier M, Pestel J, Dessaint JP, Tonnel AB, Lassalle P. Human endothelial-cell specific molecule-1 binds directly to the integrin CD11a/CD18 (LFA-1) and blocks binding to intercellular adhesion molecule-1. J Immunol. 2001;167:3099–106.

62. Scherpereel A, Depontieu F, Grigoriu B, Cavestri B, Tsicopoulos A, Gentina T, Jourdain M, Pugin J, Tonnel A-B, Lassalle P. Endocan, a new endothelial marker in human sepsis. Crit Care Med. 2006;34(2):532–7.

63. Kumar SK, Mani KP. Endocan alters nitric oxide production in endothelial cells by targeting AKT/eNOS and NFkB/iNOS signaling. Nitric Oxide. 2021;117:26–33.

64. Chen X, Zhu C, Ji C, Zhao Y, Zhang C, Chen F, Gao C, Zhu J, Qian L, Guo X. STEAP4, a gene associated with insulin sensitivity, is regulated by several adipokines in human adipocytes. Int J Mol Med. 2010;25:361–7.

65. Ohgami RS, Campagna DR, McDonald A, Fleming MD. The STEAP proteins are metalloreductases. Blood. 2006;108:1388–94.

66. Wang SB, Lei T, Zhou LL, Zheng HL, Zeng CP, Liu N, Yang ZQ, Chen X. Functional analysis and transcriptional regulation of porcine six transmembrane epithelial antigen of prostate 4 (STEAP4) gene and its novel variant in hepatocytes. Int J Biochem Cell Biol. 2013;45:612–20.

67. Chuang CT, Guh JY, Lu CY, Wang YT, Chen HC, Chuang LY. STEAP4 attenuates high glucose and S100B-induced effects in mesangial cells. J Cell Mol Med. 2015;19:1234–44.

68. Tanaka Y, Matsumoto I, Iwanami K, Inoue A, Minami R, Umeda N, Kanamori A, Ochiai N, Miyazawa K, Sugihara M, Hayashi T, Goto D, Ito S, Sumida T. Six-transmembrane epithelial antigen of prostate4 (STEAP4) is a tumor necrosis factor alpha-induced protein that regulates IL-6, IL-8, and cell proliferation in synovium from patients with rheumatoid arthritis. Mod Rheumatol. 2012;22:128-36.

69. Jin ZB, Huang XF, Lv JN, Xiang L, Li DQ, Chen J, Huang C, Wu J, Lu F, Qu J. SLC7A14 linked to autosomal recessive retinitis pigmentosa. Nat Commun. 2014;5:3517.

70. Zhuang YY, Xiang L, Wen XR, Shen RJ, Zhao N, Zheng SS, Han RY, Qu J, Lu F, Jin ZB. Slc7a14 Is Indispensable in Zebrafish Retinas. Front Cell Dev Biol. 2019;7:333.

